# A critical role for touch neurons in a skin-brain pathway for stress resilience

**DOI:** 10.1101/2022.05.23.493062

**Authors:** Melanie D Schaffler, Micah Johnson, Ben Hing, Paul Kahler, Ian Hultman, Sanvesh Srivastava, Justin Arnold, Julie N Blendy, Rainbo Hultman, Ishmail Abdus-Saboor

## Abstract

Social touch can act as a stress buffer, reducing behavioral and physiological responses to stressful scenarios. However, skin-brain touch pathways that promote stress resilience remain unknown. Here, we show that mice with an early life genetic ablation of Mrgprb4-lineage touch neurons display stress vulnerability behaviors in adulthood. Chemogenetic activation of these touch neurons reduced corticosterone levels under mild acute stress conditions. In addition, whole-brain c-Fos activity mapping while chemogenetically turning on these neurons uncovered differential neural activity patterns in brain areas relevant to somatosensation, reward, and affect. To gain mechanistic insight into this skin-brain touch pathway for stress susceptibility, we used multi-circuit neurophysiological recordings across seven brain regions at baseline and after stress in mice that had Mrgprb4-lineage touch neurons ablated in early life. Interestingly, the Mrgprb4-lineage neuron-ablated mice have alterations in local field potential phase directionality and power in the theta frequencies in mesolimbic reward regions, which may underlie our observed stress susceptibility phenotype. Together, these studies revealed that sensory neurons in the skin engage networks across the brain to promote stress resilience.

## Introduction

Tactile interaction between social mammals is a natural reward ^1–3^, motivating the formation of social bonds. Touch can also be an anxiolytic, with the ability to protect against the neurobiological and behavioral effects of stress ^4–9^. Clinical studies demonstrate that touch-based therapies like massage can reduce stress and anxiety ^10,11^ and peripherally restricted drugs targeting tactile afferents improved anxiety-like behaviors in mice ^12^. Both physical touch and drug-mediated manipulation of peripheral neurons are therefore proven to produce positive behavioral change. Analogously, a lack of social touch can result in behavioral and hormonal changes akin to anxiety and depressive disorders ^13–17^. However, the neuronal circuits from skin to brain, by which social touch, or the lack thereof, can alter mood remains unknown. Here, we aim to identify the peripheral sensory neurons in the skin of mice that are at the start of such a circuit.

Social touch behaviors like hugging, huddling, and social-grooming activate a wide variety of sensory neurons, but only a subset govern the pleasant nature of touch. Here, we focus on Mas-related G-protein coupled receptor B4 (Mrgprb4) -lineage sensory neurons in mice. Mrgprb4+ touch neurons in mice respond to touch stimuli, and animals develop a preference for being in a chamber where Mrgprb4 neurons were previously chemogenetically activated ^18^. These neurons also project exclusively to the hairy skin (areas predominantly involved in social touch behaviors like allo-grooming) ^19^, and they provide input to an affective touch spinoparabrachial pathway ^20^. In our recent studies, we demonstrate that optogenetic activation of Mrgprb4-lineage touch neurons stimulates dopamine release in the nucleus accumbens and promotes conditioned place preference^21^. Thus, these neurons appear to be a part of a rewarding touch circuit in mice, but it remains unknown whether these neurons might also play a role in the calming, anxiolytic effects of social touch.

In the present study, we sought to determine (1) the long-term effects of early-life ablation of Mrgprb4 lineage touch neurons on behavior and neurophysiology and (2) the rescue potential of Mrgprb4-lineage neuron activation on stress-induced behaviors and hormonal changes in adulthood. Based on the importance of Mrgprb4-lineage neurons in social reward^22^ and the emotional consequences of touch or the lack thereof across species, we hypothesized that the developmental ablation of Mrgprb4-lineage neurons would increase the stress response in adulthood, and that their activation would be therapeutic during stress. Our findings indicate that Mrgprb4-lineage neurons play a critical role in stress resilience and that their activation can reduce the endocrine response to mild stress. Also, neurophysiological recordings across seven distinct brain regions previously implicated in reward and stress susceptibility suggest a potential mechanism that underlies this stress susceptibility.

## Results

### Mice with Mrgprb4-lineage neurons ablated display normal cognitive, locomotor, discriminative touch, and mood-related behaviors

To determine whether the loss of a subset of touch neurons implicated in social touch would impact development or adult behavior, we crossed *Mrprb4*^*Cr*e^ mice to *Rosa*^*DTA*^ mice to genetically ablate Mrgprb4 neurons (Fig1A). Because Mrgprb4 expression begins around postnatal day (PND 2) ^19^, and the cre-dependent ablation method ablates cells ∼1 day after Cre expression starts ^23^, Mrgprb4 lineage neurons were likely ablated ∼PND 3. Using RNAscope *in situ* hybridization, we confirmed the extinction of cells that express Mrgprb4 (Fig 1B-C). Because Mrgprb4 is expressed in the dorsal root ganglion more broadly at the PND 3 timepoint than in adulthood, populations of sensory neurons that share a developmental history, including a fraction of the Mrgprd+ and Mrgpra3+ neurons, are also ablated (Fig 1C). We therefore refer to our manipulated population as Mrgprb4-lineage neurons. These Mrgprb4-lineage neuron-ablated mice did not display any developmental (Fig 1D-I), cognitive (*F*_(1,24)_ = 1.622, *p* = 0.8715; Fig 1J), locomotor (*F*_(1,27)_ = 1.047, *p* = 0.5108; Fig 1K), or discriminative touch deficits (*F*_(1,14)_ = 1.730, *p* = 0.5202; Fig 1L) that could confound other behavioral results.

**Figure 1.**
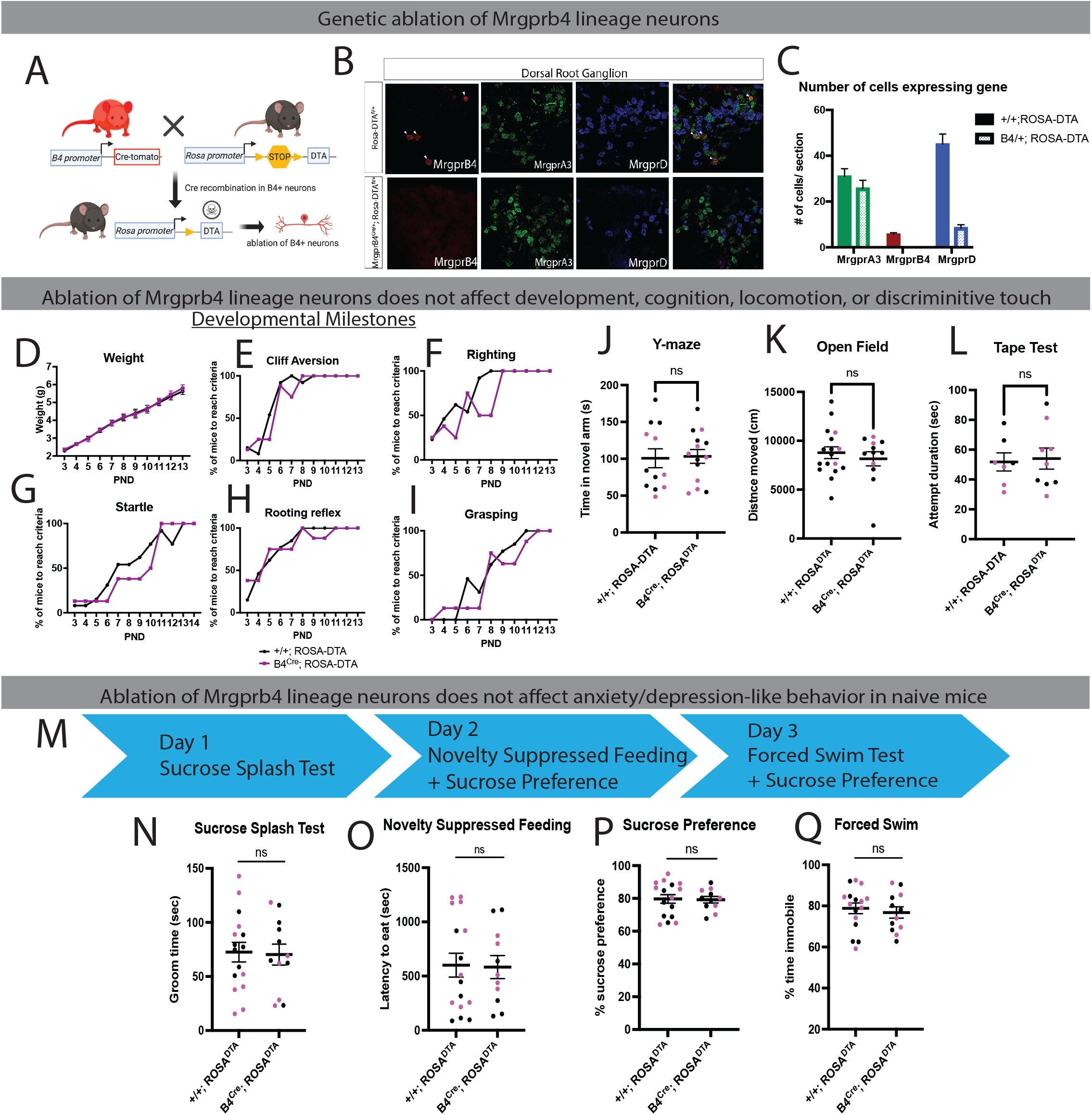
Ablation of Mrgprb4 lineage neurons does not affect development or anxiety/depression -related behavior at baseline. **A**, Generation of Mrgprb4^Cre^; Rosa^DTA^ mice. **B**, RNA Scope *in situ* hybridization in DRG to quantify the ablation of Mrgprb4 lineage neurons. **C**, Quantification of ablation of different populations of DRG neurons that share lineage with Mrgprb4 (n=4 mice/group, 6 sections/mouse). **D-I**, Mice with Mrgprb4 lineage neurons ablated do not display developmental deficits (n=8-13/group). **J**, There was no effect of Mrgprb4 lineage ablation on time spent in novel arm in the Y-maze assay. **K**, Ablation did not affect locomotor activity in an open field. **L**, Ablation did not affect discriminative touch ability. **M**, Anxiety/depression-related behavioral testing schedule. **N**, Ablation did not affect groom time in sucrose splash test. **O**, Ablation did not affect latency to eat in the Novelty Suppressed Feeding Assay. **P**, Ablation did not affect preference for sucrose. **Q**, Ablation did not affect time immobile in the Forced Swim Test. Data were analyzed by unpaired t-test (n=12-16/group). Magenta circles indicate female mice.

To determine if the ablation of Mrgprb4-lineage neurons impacts circuits related to negative affect, we tested the mice in the sucrose splash test (*F*_(1,26)_ = 1.186, *p* = 0.8701; Fig 1N), novelty suppressed feeding (NSF) (*F*_(1,25)_ = 1.574, *p* = 0.9128; Fig 1O), sucrose preference (*F*_(1,25)_ = 2.499, *p* = 0.8970; Fig 1P), and forced swim (FST) (*F*_(1,26)_ = 1.186, *p* = 0.6050; Fig 1Q) tests. These assays are commonly used assessments of rodent self-care behavior and motivation for reward, fear of novel spaces, anhedonia, and helplessness behavior, respectively, and serve as proxies for neural circuits whose activity is linked to underlying domains of anxiety/depression. *Mrgprb4*^*Cre*^; *Rosa*^*DTA*^ males and females behaved similarly to controls on all 4 behavioral assays, suggesting that the loss of Mrgprb4 lineage neurons does not impact anxiety/depression-related behavior in mice at baseline. Overall, ablation of this relatively small population of peripheral sensory neurons is not sufficient to alter development or lead to gross behavioral deficits.

### Mice with Mrgprb4-lineage neurons ablated display an anxiety/depression-like phenotype after stress

Early-life stress in humans can have long lasting effects on affective and cognitive function and is associated with increased risk of depression^24,25^. Similarly in mice, stress or a genetic manipulation during development can increase susceptibility to adult stress ^26–28^. We therefore tested whether *Mrgprb4*^*Cre*^; *Rosa*^*DTA*^ mice would behave similarly to controls after stress in adulthood. We also collected blood to assess any differences in circulating corticostone levels as a proxy for hypothalamic pituitary adrenal (HPA) axis activity. Before exposure to the same anxiety/depression assays used in Figure 1, mice were exposed to 3 days of variable stress, which is a previously reported subthreshold variable stress paradigm ^27^. Blood was collected during a brief restraint the day before the variable stress began, 30 minutes after FST, and 24h after FST (Fig 2A), and CORT levels were measured. Similar to testing under pre-stress conditions (Fig 1), Mrgprb4-lineage ablation did not affect behavior in the novelty suppressed feeding (*F*_(1,30)_ = 1.123, *p* = 0.1629; Fig 2C) or sucrose preference assays (*F*_(1,19)_ = 2.421, *p* = 0.2100; Fig 2D).

**Figure 2.**
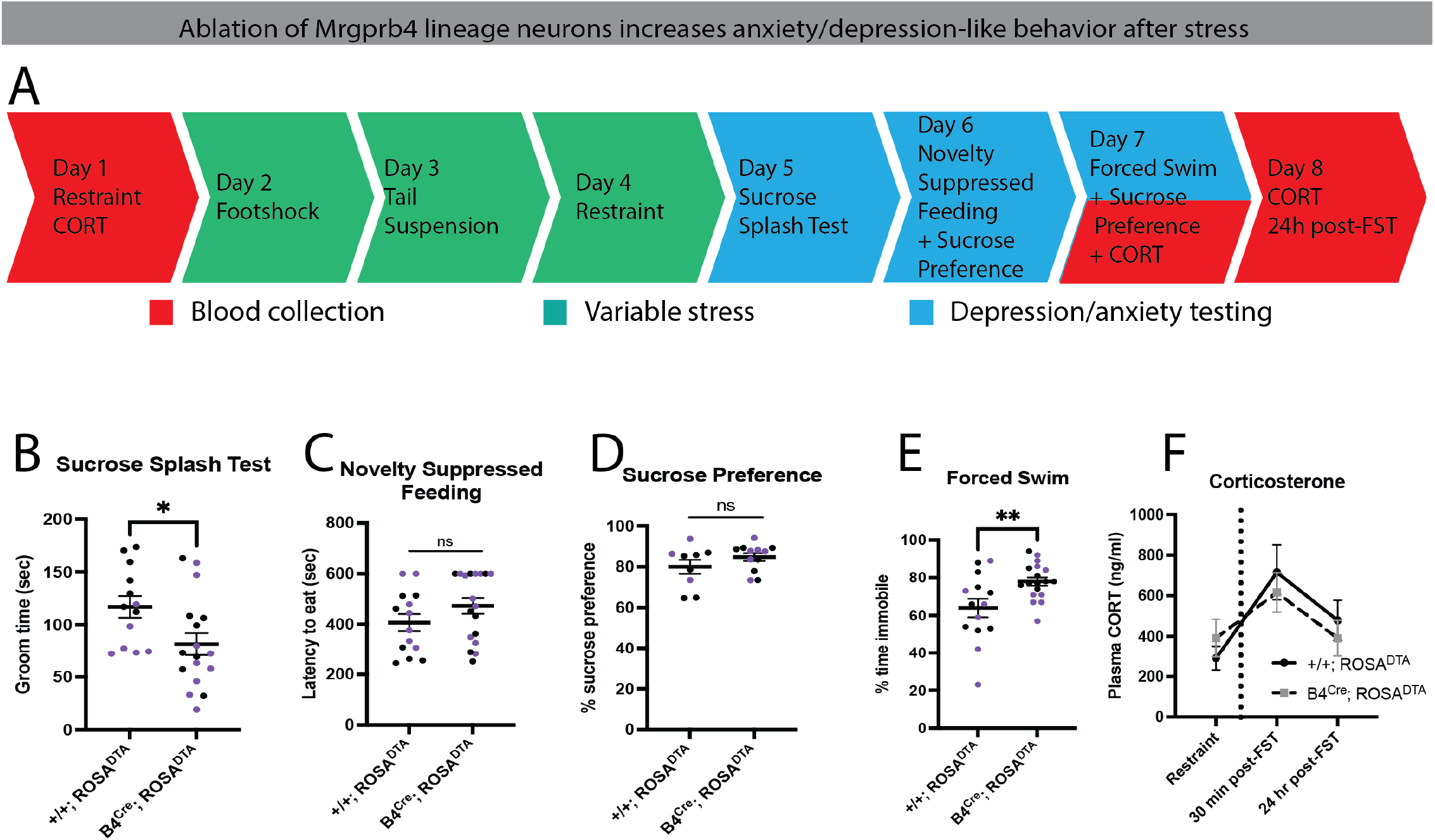
Ablation of Mrgprb4-lineage neurons increases anxiety/depression-like behavior after stress. **A**, Testing schedule for blood collection, variable stress, and behavioral testing. **B**, Mice with Mrgprb4 lineage neurons ablated had decreased groom time in the sucrose splash test. **C**, Ablation did not affect latency to eat in the Novelty Suppressed Feeding Assay. **D**, Ablation did not affect preference for sucrose. **E**, Ablation of Mrgprb4 lineage neurons increased immobile time in the forced swim test. **F**, Mice with Mrgprb4 lineage neurons ablated have a similar corticosterone response to FST stress as controls. Data were analyzed by unpaired t-test (n=14-18/group). Purple circles indicate female mice.

Interestingly, we found that stressed mice with Mrgprb4-lineage neurons ablated spent less time grooming in the sucrose splash test, indicating decreased self-care behavior and potentially a reduced motivation for reward (*F*_(1,28)_ = 1.397, *p* = 0.0259; Fig. 2B). These Mrgprb4-lineage neuron ablated mice also had increased immobile time in the forced swim test, indicating increased helplessness in response to an aversive stimulus (*F*_(1,30)_ = 3.743, *p* = 0.0086; Fig. 2E) relative to controls. Further, as expected, we observed a pattern in which the corticosterone levels in control mice spiked 30 minutes after forced swim and fell 24 hours later, though not back to pre-variable stress levels. Despite a behavioral difference between groups in the forced swim test, both groups displayed a similar pattern in corticosterone levels, demonstrating a disconnect between behavior and endocrine hormone levels. Taken together, these findings uncover a role for Mrgprb4-lineage neurons in promoting behavioral resilience following physical stressors.

### Activation of Mrgprb4-lineage neurons reduces corticosterone levels under mild stress conditions but does not rescue anxiety/depression-related behavior after variable stress

After identifying several behavioral stress susceptibility phenotypes after early life ablation of Mrgprb4-lineage neurons, we next asked if chemogenetic activation of these neurons in adulthood during stress induction was protective. To test this, we subjected *Mrgprb4*^*Cre*^; *Rosa*^*hM3dq*^ and control mice to a testing schedule like that used in the ablation experiments. Mice were given CNO (1mg/kg) each day, 2 hours before the start of the stressors and behavioral experiments (Fig 3C). This time interval was chosen because 2 hours marks the end of the CNO peak efficacy window ^29^. Our goal here was to potentially mimic the calming, social touch that naturally occurs between cage mates. We therefore wanted the neurons to be active while the mice were together in the home cage, in a social context, and not during the behavioral tests.

**Figure 3.**
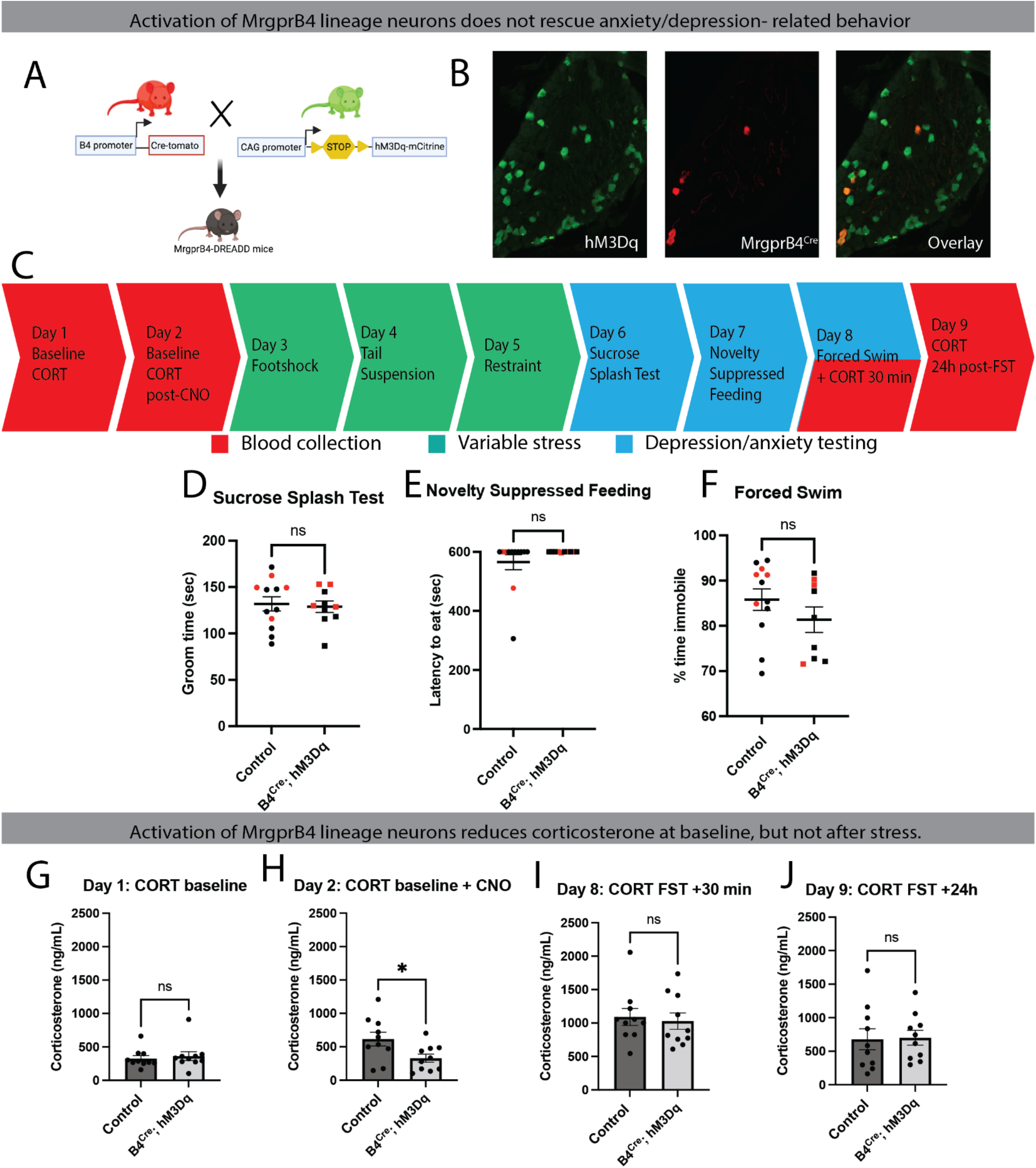
Activation of Mrgprb4 lineage neurons with DREADDs reduces corticosterone after an acute mild stress but does impact behavior after variable stress. **A**, Generation of *Mrgprb4*^*Cre*^; *Rosa*^*hM3dq*^ mice. Control mice are *Mrgprb4*^*Cre*^; *+/+ and +/+; Rosa*^*hM3dq*^ littermates. **B**, Immunohistochemistry in the DRG of *Mrgprb4*^*Cre*^; *Rosa*^*hM3dq*^ mice to confirm presence of hM3dq receptors in Mrgprb4+neurons. **C**, Testing schedule for blood collection, variable stress, and behavioral assays. **D**, Activation did not alter behavior in the sucrose splash test. **E**, Activation did not alter behavior in the novelty suppressed feeding assay. **F**, Activation does not impact behavior in the forced swim test. **G**, *Mrgprb4*^*Cre*^; *Rosa*^*hM3dq*^ mice and controls have similar CORT levels at baseline. **H**, *Mrgprb4*^*Cre*^; *Rosa*^*hM3dq*^ mice and controls have similar CORT levels after mild acute stress. **I**, *Mrgprb4*^*Cre*^; *Rosa*^*hM3dq*^ mice have reduced CORT levels after CNO injection and mild acute stress. **J**, *Mrgprb4*^*Cre*^; *Rosa*^*hM3dq*^ mice and controls have similar CORT levels 30min after FST. **K**, *Mrgprb4*^*Cre*^; *Rosa*^*hM3dq*^ mice and controls have similar CORT levels 24h after FST. Data were analyzed by unpaired t-test (n=10-12/group). Red circles indicate female mice.

The presence of hM3dq receptors in Mrgprb4 neurons was confirmed using immunohistochemistry (Fig 3B). Though 100% of Mrgprb4 neurons expressed hM3Dq receptors, Mrgprb4 neurons represented only ∼10% of all neurons expressing the DREADDs. Based on the quantification of cre-dependent neuronal ablation (Fig 1C), and the previous work characterizing the expression of cre-dependent strategies in these mice, the remaining ∼90% are likely a subset of Mrgprd+ and MrgprA3+ neurons. This is consistent with the expected developmental expression of Mrgprb4 ^30^. We believe the co-expression of hM3dq in Mrgprd+ and Mrgpra3+ in this experiment is tolerable since we do not label the entire populations of these neurons, and moreover, we do not observe any pain or itch related phenotypes after CNO injection, which are sensations that these neurons encode.

Activation of Mrgprb4-lineage neurons did not prevent stress-induced behaviors in the sucrose splash test (*F* _(1,20)_ = 1.809, *p* = 0.7647; Fig 3D), novelty suppressed feeding assay (*F* _(1,20)_ = 8795, *p* = 0.2369; Fig 3E), or forced swim test (*F*_(1,19)_ = 1.067, *p* = 0.2431; Fig 3F). We did observe a reduction in corticosterone levels after mild acute stress with CNO injection on Day 3 of the testing schedule, prior to the 3 days of variable stress (*F* _(1,18)_ = 2.810, *p* = 0.0285; Fig 3I). We did not observe a reduction 30 minutes (*F* _(1,18)_ = 6.766, *p* = 0.2898; Fig 3J) or 24h (*F* _(1,18)_ = 1.495, *p* = 0.8453; Fig 3K) after the forced swim test. Therefore, activation of Mrgprb4-lineage neurons may be able to reduce the stress response after an acute, mild stressor but not after more severe, repeated stress.

### DREADD-mediated activation of Mrgprb4-lineage neurons alters activity brain areas relevant to somatosensation and affect

Reductions in goal-directed behaviors, like those observed in Figure 2, have been attributed to multiple brain regions, including the amygdala, ventral hippocampus, nucleus accumbens, and ventral tegmental area^31–33^. It is therefore interesting that we observe such behavioral differences after manipulating neurons in the skin. Thus, we aimed to gain insight into how Mrgprb4-lineage neurons may be connected to circuits in the brain. To assess the downstream effects of Mrgprb4-lineage neuron activation on the brain, we used SHIELD-based tissue phenotyping (Fig 4B). Stress-naïve Mrgprb4^Cre^; Rosa^hM3dq^ mice and controls were both given CNO (Fig 4A) and perfused 2 hours later to assess peak c-Fos activity. Brains were cleared, immunolabled for CFos and NeuN, and imaged using light sheet microscopy. The images were then aligned to the Allen Brain Atlas and cell counts were determined using automated cell detected software.

**Figure 4.**
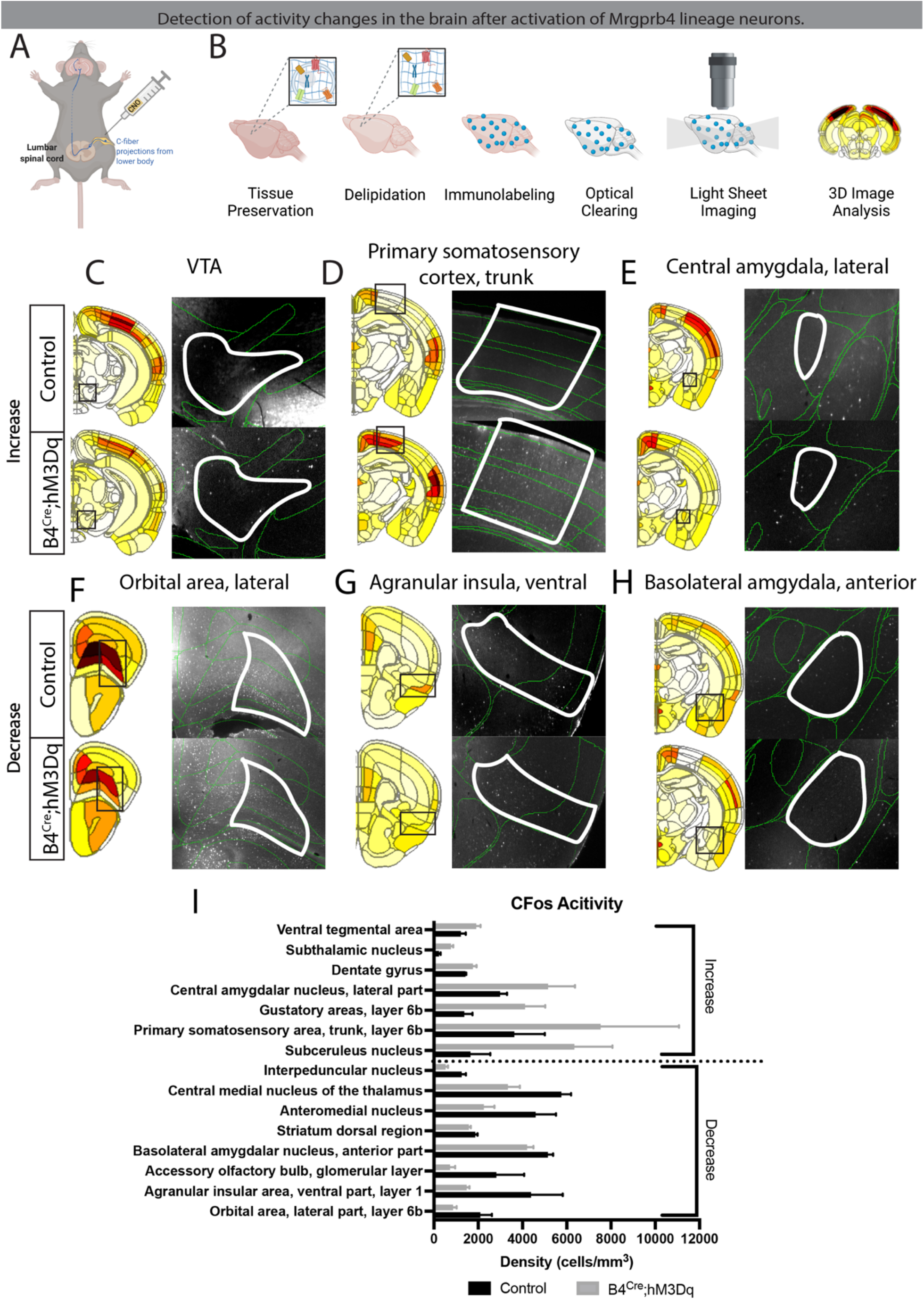
Activation of Mrgprb4-lineage neurons impacts brain activity in sensory and affect-related brain areas. **A**, All mice received CNO via IP injection. **B**, Brain tissue processing steps. **C**, Activity in the VTA increased with Mrgprb4 lineage neuron activation. D, Activity in the primary somatosensory cortex increased with Mrgprb4 lineage neuron activation. **E**, Activity in the lateral central amygdala increased with Mrgprb4 lineage neuron activation. **F**, Activity in the lateral orbital areas decreased with Mrgprb4 lineage neuron activation. **G**, Activity in the ventral agranular insular areas decreased with Mrgprb4 lineage neuron activation. **H**, Activity in the anterior basolateral amygdala decreased with Mrgprb4 lineage neuron activation. **I**, Cfos+ cell density in select brain regions.

Among the most robustly activated brain regions were the ventral tegmental area (VTA), primary somatosensory cortex, and the lateral part of the central amygdala (CeAl) (Fig 4C-E). Importantly, the primary somatosensory cortex, especially the portion receiving input from the body’s trunk, as it is comprised completely of hairy skin, was activated upon stimulation of Mrgprb4-lineage neurons. This activation pattern indicates that we are in fact activating a somatosensory skin-to-brain circuit. Further, neurons in the VTA play a critical role in reward behavior ^34^ and the lateral subdivision of the CeA controls responses to aversive stimuli by inhibiting the output of the medial CeA ^35^, the amygdala’s primary output region ^36^.

Among the areas with the largest decrease in activity were the lateral orbital area, ventral part of the agranular insula, and the anterior part of the basolateral amygdala (BLA) (Fig 4F-H). In rodents, the lateral orbital area is a subdivision of the orbitofrontal cortex (OFC), an area important for emotional processing. Chronic inactivation of the lateral orbital area reduced depression-like behavior in the forced swim test in rats ^37^, indicating that a reduction of activity in this area can improve negative valence. Additionally, rodent projection neurons that originate in the ventral agranular insular cortex terminate in the infralimbic and prelimbic areas^38^, two areas important in the fear response ^39^. Finally, the basolateral amygdala is divided into an anterior and a posterior part, both of which project directly to ventral hippocampal CA1. Activation of the anterior BLA-vCA1 inputs induces anxiety and social deficits ^40^, suggesting that a *reduction* of activity in anterior BLA may have anxiolytic effects. Overall, the activation of Mrgprb4-lineage neurons alters brain activity in multiple brain areas involved in anxiety, fear, emotional processing, and somatosensation. These experiments provide a comprehensive map of brain activity upon Mrgprb4-lineage activation and provide a basis for future to studies to look deeper at cell type specificity and directionality in brain networks with Mrgprb4-lineage neuron activation.

### Mice with Mrgprb4-lineage neurons ablated have altered LFP signal directionality in valence-relevant brain areas at baseline

To gain insight into this skin-brain touch pathway for stress susceptibility, guided by our c-Fos results and previous work discovering spatiotemporal dynamic networks that predict stress vulnerability, we used multi-circuit neurophysiological recordings in the infralimbic cortex (IL), prelimbic cortex (PrL), nucelus accumbens (Nac), ventral hippocampus (vHipp), ventral tegmental area (VTA), basolateral amygdala (BLA), and central amygdala (CeA) at baseline in *Mrgprb4*^*Cre*^; *Rosa*^*DTA*^ mice and controls^41^ (Fig5 A,B). This multi-site in vivo recording approach quantifies local field potentials (LFPs). Correct electrode tip placement in each of the seven brain regions was verified (Supplemental Fig1). We used a phase offset approach to estimate signal directionality between each pair of brain regions for both *Mrgprb4*^*Cre*^; *Rosa*^*DTA*^ mice and controls at baseline conditions. This measurement provides an estimate of information transfer based on quantifying the amount of time by which one of two synchronous LFPs leads the other^41–43^. Such a strategy has been used successfully to identify frequency bands of biological and behavioral relevance^44,45^. Because we have simultaneous data from seven recording sites, we identified signal directionality for 21 brain region pairs (Fig. 5, Supplemental Fig2). We used a lasso-based method to specifically identify such frequency bands of interest (Supplemental Fig3). Similarly to previous studies, directionality between PFC regions (PrL and IL) and AMY regions (BLA and CeA) identified PFC regions leading in the 2-7Hz frequency band and AMY regions leading in the 8-11Hz range. Interestingly, NAc-VTA directionality was strikingly different between genotypes. At 1-8Hz, the NAc was leading by ∼10-15ms +/-3ms in both genotypes. At 9-10Hz, the VTA is leading by 41ms +/-5.5ms in control mice but by only 16ms+/-2.6ms in *Mrgprb4*^*Cre*^; *Rosa*^*DTA*^ mice (Fig5 C-F). Given this interesting difference, and our prior findings connecting Mrgprb4-lineage neurons to brain reward areas^22^, we decided to direct our attention to the VTA-Nac circuit. The dopaminergic inputs from VTA to NAc drive reward signaling^46,47^ and are altered in *Mrgprb4*^*Cre*^; *Rosa*^*DTA*^ mice (Fig5F). These results suggest that an alteration in reward processing is linked to the stress susceptibility phenotypes.

**Figure 5.**
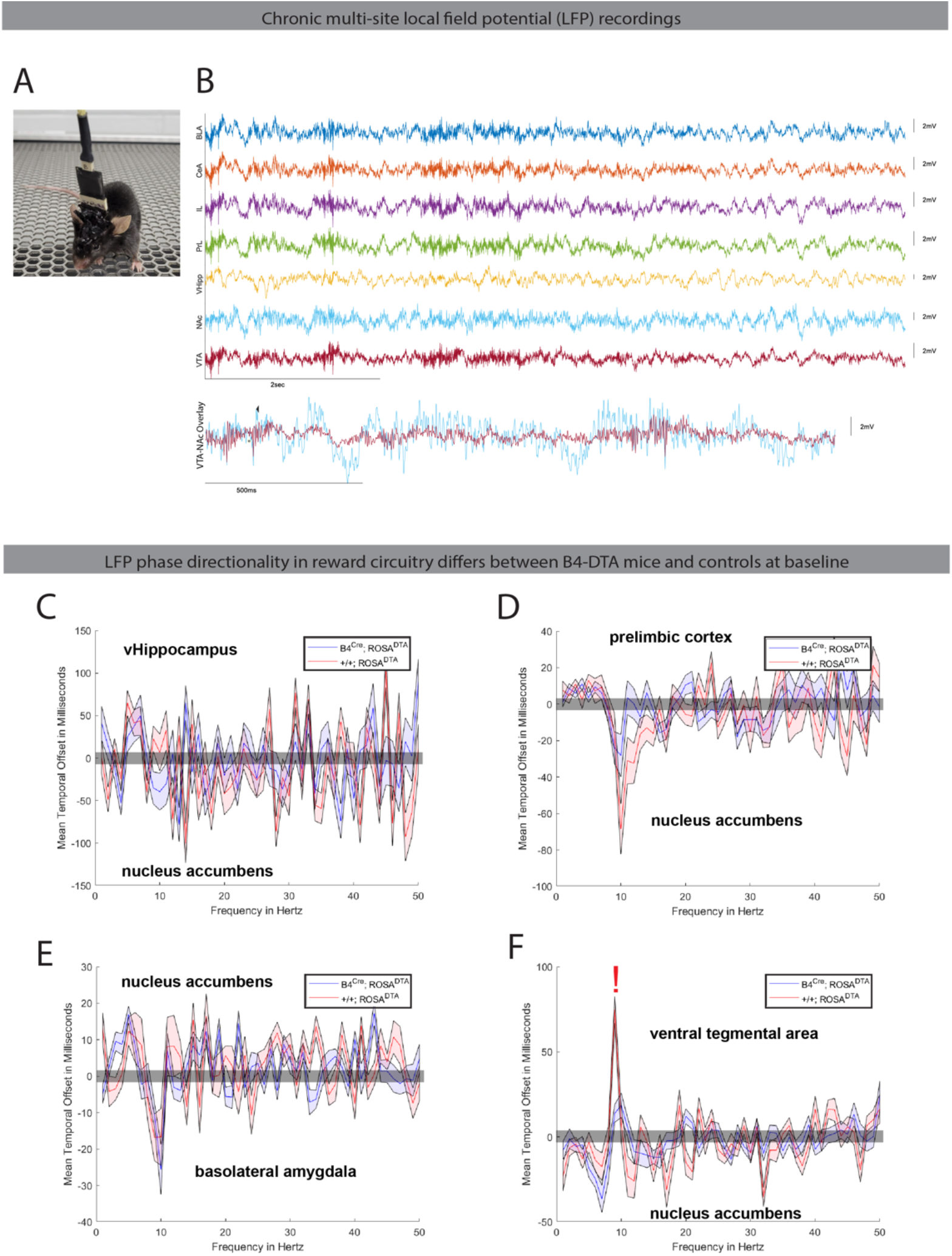
Ablation of Mrgprb4-lineage neurons alters LFP signal directionality in the VTA-Nac circuit in stress-naïve mice. **A**, Mice were implanted with electrodes in 7 different brain regions. **B**, Electrodes were placed into the VTA, Nac, Vhipp, PLC, IL, CeA, and BLA. Representative LFP traces from a control animal. **C**, The phase offset time series was calculated between the vHipp and Nac, **D**, the PLC and Nac, **E**, the Nac and BLA, and **F**, the VTA and Nac. The red “!” indicates the frequency at which the VTA leads the NAc to a larger degree in control mice.

### Mice with Mrgprb4-lineage neurons ablated display altered brain activity in the NAc after stress

Lower frequency changes in phase directionality between the VTA and NAc were observed during baseline conditions (Fig 5G). We therefore next sought to test if oscillatory power in these reward-related regions was altered after stress in the frequency band in which VTA is leading (9-10Hz) and in the frequency band in which NAc is leading (1-8Hz). These recordings were performed post-variable stress, during the sucrose splash test, where we previously observed stress-induced reductions in grooming behavior in *Mrgprb4*^*Cre*^; *Rosa*^*DTA*^ mice (Figure 2B). After the 3-day variable stress paradigm, LFPs were recorded in an empty home cage environment for 5 minutes. After the 5min, mice were then sprayed with sucrose solution and LFPs were recorded for an additional 5 mins. During the sucrose portion of the test, NAc 1-8 Hz power was significantly reduced in *Mrgprb4*^*Cre*^; *Rosa*^*DTA*^ mice (Fig6E). Both before and after being sprayed with sucrose solution, *Mrgprb4*^*Cre*^; *Rosa*^*DTA*^ mice exhibited lower LFP power in the NAc, in the lower theta frequency range (Fig 6C-E). Interestingly *Mrgprb4*^*Cre*^; *Rosa*^*DTA*^ mice with the lowest NAc power in the theta frequency range, also displayed the strongest behavioral deficit in the sucrose test after stress (Fig 6D,F). Together, these findings suggest that reduced neurophysiological power in the NAc might be causally related to the stress susceptibility behavior revealed when Mrgprb4-lineage neurons are ablated.

**Figure 6.**
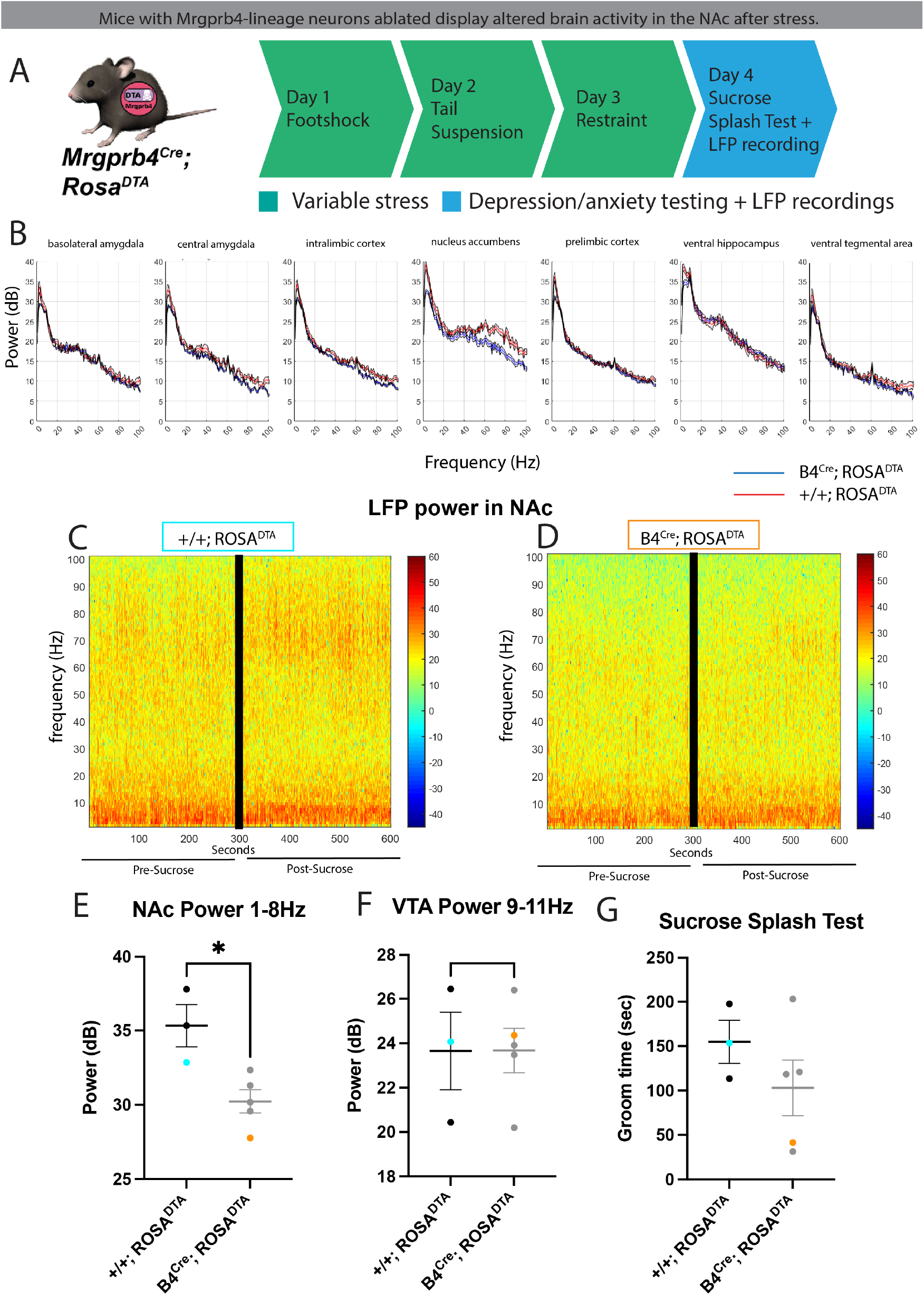
Ablation of Mrgprb4-lineage neurons alters brain activity in the NAc after stress. **A**, Representation of *Mrgprb4*^*Cre*^; *Rosa*^*DTA*^ mice and behavioral schedule utilized. **B**, B4^Cre^; Rosa^DTA^ and +/+;Rosa^DTA^ animals show differences in nucleus accumbens power after subchronic variable stress during the sucrose splash test + paired recordings **C**, Representative power spectrograms in the nucleus accumbens of +/+;Rosa^DTA^ and **D**, B4^Cre^; Rosa^DTA^ animals. **E**, B4^Cre^; Rosa^DTA^ mice had decreased NAc power in the 1-8 Hz frequency range. **F**, +/+;Rosa^DTA^ and B4^Ce^; Rosa^DTA^ mice have similar VTA power in the 9-11Hz range. **G**, Mice with ablated Mrgprb4-lineage neurons spent less time grooming than +/+;Rosa^DTA^ mice during the sucrose splash test + paired recordings. Data were analyzed by unpaired t-test (n=3-5/group). Light blue dots indicate representative control animal and orange dots indicate representative B4^Cre^; Rosa^DTA^ animal.

## Discussion

The present study demonstrates that manipulation of peripheral Mrgprb4-lineage neurons can alter behavior, stress hormone levels, and brain activity. Developmental ablation of Mrgprb4-lineage neurons increased depression-like behavior after variable stress. DREADD-mediated activation of Mrgprb4 lineage neurons in adulthood was not sufficient to rescue or prevent the effects of stress on behavior, but did reduce corticosterone levels after mild acute stress. Neurophysiological coupling between the VTA and NAc, a key pathway involved in reward and prevention of stress-induced depression, was altered before and after stress when Mrgprb4-lineage neurons were ablated. Together, these data provide evidence that Mrgprb4-lineage neurons are important for normal stress resilience in adulthood, play an important role in the long-term benefits of touch during development, and might have therapeutic potential if an ideal activation regimen is identified.

### A novel early life stress model

The ablation of Mrgprb4-lineage neurons at ∼PND 3 resulted in increased susceptibility to stress in adulthood. Maternal separation and deprivation paradigms are commonly used, well-validated depression models ^48,49^. However, in these paradigms, pups are deprived of food and warmth and placed in an unfamiliar space, in addition to the loss of touch. We were therefore previously unable to assess which component of maternal separation causes the long-term negative consequences. The present study suggests that simply altering the tactile experience of mice by ablating a subset of mechanosensitive neurons, one that does not impact general discriminative touch (Fig 1L), is sufficient to induce long-term behavioral change. This work introduces a new early-life stress model in which the home cage and dam can remain largely undisturbed, with minimal handling and without disrupting pups’ access to food, warmth, and cage mates. This could be beneficial for studies with severely immunocompromised or otherwise sensitive mice in which mouse handling and mother manipulations need to be restricted.

### Developmental vs adult manipulation of Mrgprb4-lineage neurons

We demonstrate that ablation of Mrgprb4-lineage neurons at ∼PND 3 reduces resilience to stress in adulthood. We also tested whether specific activation of MrgprB4+ neurons can promote stress resilience, as measured by behavioral outcomes and corticosterone levels. Based on the results of the ablation experiments, and previous research demonstrating that tactile stimulation can rescue stress-induced behaviors ^4,50,51^, we hypothesized that activating Mrgprb4-lineage neurons would be sufficient to rescue the behavioral phenotypes observed after stress and that corticosterone levels would mirror this. However, our results suggest that these neurons do not have therapeutic potential under the stress conditions used. These data might point toward presence of a critical window for manipulation of these neurons. Perhaps, if we activated these neurons during development and throughout life, we would observe a behavioral or more robust endocrine change.

### Relevance of neuronal activation technique

For our Mrgprb4-lineage neuron activation experiments, we used DREADDs to test whether increased activity of these neurons prior to stress was protective. Previous work from the lab found that optogenetic activation of these neurons promotes conditioned place preference and stimulates dopamine release from the nucleus accumbens. For these experiments, we chose a chemogenetic approach to look at the effects on activation in a natural, social context rather than flashing multiple pulsing lasers of a visible wavelength into the home cage. However, it may be that the chemogenetic activation of these neurons is not as rewarding. The optogenetic approach produces a smaller, more focal activation of only the back skin which more closely mimics natural behaviors like allo-grooming.

Additionally, there are multisensory contributions to the pleasantness of affective touch. Touch sensation is modulated by auditory ^52,53^ olfactory ^54,55^, and visual ^56^ cues. Perhaps because these mice were placed in a temporary holding room during Mrgprb4 lineage neuron activation, their perception of the sensation was affected. Maybe repeating the same experiments while the mice were kept in the housing room where they spent most of their lives would produce different behavioral outcomes. The unfamiliar context could have competed with the Mrgprb4-lineage neuron stimulation.

### Relationship to human CT afferents

It is tempting to speculate that the data here suggests that an analogous population of neurons in humans would play a similar role in soothing touch. Mrgprb4 neurons in mice project exclusively to the hairy skin, are unmyelinated, and have free nerve endings with large terminal arborizations ^19^. These unique anatomical features are reminiscent of C-tactile (CT) afferents in humans, a subset of neurons purported to detect pleasant, social touch^57–60^. In addition to anatomical features, CT afferents in humans and Mrgprb4+ neurons in mice also share functional similarities, as they both respond to gentle stroking ^18,19,57^, and convey an affective quality of touch^18,21^ and CT afferent firing rates positively correlate with rating of touch “pleasantness” ^58,61^.

However, the association between CT afferents and affective touch in humans is correlative, and the C-tactile hypothesis is still debated. Further, evidence that Mrgprb4+ neurons are homologous to CT afferents is sparse, and due to the nature of our genetic targeting strategy of the Mrgprb4-lineage, as opposed to the MrgprB4 adult population, this is not a pure C-low threshold mechanosensory neuron population. Regardless of whether MrgprB4+ neurons, or the Mrgprb4-lineage neurons, are analogous or related to human CT afferents, the results of this study indicate that pharmacological activation of peripheral sensory neurons can alter activity in affect-related brain areas and has the ability to modulate the stress response in mammals.

### Elucidating the skin-to-brain soothing touch circuit

Our data indicate that both developmental ablation and chemogenetic activation of Mrgprb4 lineage neurons in the skin modulate activity in the brain. The developmental ablation of Mrgprb4-lineage neurons altered spatiotemporal dynamic networks, particularly in brain reward areas, in adulthood. DREADD-mediated activation of these peripheral neurons altered activity in brain areas linked to reward and emotional processing. These two sets of experiments, congruent with previous work from the lab^22^, suggest that Mrgprb4-lineage neurons act on affect-relevant brain circuits.

Neurophysiological recordings demonstrated a reduction in LFP power in the NAc in Mrgprb4-lineage neuron-ablated mice during post-stress behavior. Moreover, Mrgprb4-lineage neuron ablated mice have altered signal directionality between the VTA and NAc, even at baseline before stress induction.

These findings mirror recent results showing that mice with higher activity in D1 medium spiny neurons in the NAc, are more resistant to subsequent stress^62^. Further, the LFP oscillatory power was observed to be significantly different in Mrgprb4-lineage ablated mice in the 1-8Hz range. This finding is intriguing given an exciting recent study that demonstrated that restraint stress alters the NAc LFP around 4Hz and that this change predicts subsequent reduction of reward seeking^63^. Thus, our results suggest that a reduction in NAc LFP power, and a lack of normal time offset phase directionality between the VTA and NAc may preclude the *Mrgprb4*^*Cre*^; *Rosa*^*DTA*^ mice towards stress susceptibility.

In conclusion, the results of these experiments further our understanding of the relationship between the peripheral nervous system and stress. Future experiments may expand upon these findings by using viral tracing techniques to determine pathways from skin to spinal cord to brain, by which activation of MrgprB4-lineage neurons in the periphery influences the brain to alter behavioral output. It would also be interesting to test whether strategies that restore the temporal LFP offset between the VTA and NAc, which was diminished when we ablated touch neurons, could be used to rescue deficits driven by reduced early life touch. Ultimately, this study might provide a framework for using the peripheral nervous system as a therapeutic target. This could be particularly beneficial for individuals without access to social touch, like infants in the NICU or severely immunocompromised adults.

## Materials and Methods

### Animals

All procedures were approved by the Institutional Animal care and Use Committee (IACUC) of the University of Pennsylvania, Columbia University, and the University of Iowa. Male and female mice aged 8-12 weeks were used for all behavioral experiments except developmental milestones for which mice were tested PND 3-14. MrgprB4^cre^ mice were bred at the testing sites on a C57bl/6J background from a line originally generated in the laboratory of David Anderson at the California Institute of Technology ^18^. Mice were group housed and maintained on a 12 h light/dark cycle with ad libitum access to food and water, except when explicitly stated for behavioral testing. MrgprB4^Cre^; Rosa^DTA^ mice were single housed following the sucrose splash test. Mouse lines used in this study were purchased from Jackson Laboratories: *Mrgprb4*^*Cre*^ (Stock No: 021077), *Rosa*^*DTA*^ (Stock No: 009669), *Rosa*^*Gq-DREADD*^ (Stock No: 026220).

### Immunohistochemistry

Brains used for CFos mapping were harvested immediately following perfusion and post-fixed in 4% PFA at 4°C for 24h. Brains were then then placed in 0.02% sodium azide and shipped to LifeCanvas Technologies for tissue clearing, antibody labeling, imaging and analysis. DRGs used to confirm the co-expression of DREADDs and Mrgprb4^Cre^ were harvested immediately following perfusion, post-fixed in 4% PFA at 4°C for 24h, and then 30% sucrose for another 24h. Tissue was flash frozen on dry ice in OCT, sectioned at 20um onto Superfrost Plus Slides, and stored at -20 °C. IHC protocol: 3 × 5 min PBS washes; 30 min permeabilization in PBST (0.3% triton); 1 hour blocking in PBST with 5% normal donkey serum (NDS); GFP primary antibody 1:1000 and RFP antibody 1:500 in PBST with 5% NDS applied to slides in humidified chamber overnight at 4°C. Secondary antibody [1:250] in PBST with 5% NDS was applied to slides in humidified chamber for 1.5h at room temperature. Slides were washed 3 × 10 min in PBS before applying mounting media and a coverslip.

### Whole mouse brain processing, staining, and imaging

Whole mouse brains were processed following the SHIELD protocol (LifeCanvas Technologies ^64^. Briefly, SHIELD is a tissue preservation technique that enhances standard PFA fixation. This technology allows for the extraction of multiple molecules from the same transparent tissue. It is a preparation step necessary for active clearing or immunolabeling, as it protects important targets from harsher chemical or environments that can strip tissue. Samples were cleared for 1 day at 42°C with SmartBatch+ (LifeCanvas Technologies), a device employing stochastic electrotransport^65^.

Cleared samples were actively immunolabeled using SmartBatch+ (LifeCanvas Technologies) based on eFLASH technology integrating stochastic electrotransport^65^ and SWITCH ^66^. SmartBatch+ combines active tissue clearing and immunolabeling into one device. Stochastic electrotransport selectively enhances transport of highly electromobile molecules, like antibodies, through a porous sample. It speeds up the diffusion time for immunolableing, which can take a long a long time for large samples like an intact mouse brain. eFLASH is a tissue labeling technique that allows for uniform whole-organ staining in <24 hours ^67^and SWITCH controls chemical reactions during tissue processing to ensure uniformity of preservation and immunolabeling^66^. Each brain sample was stained with primaries, 3.5 mg of rabbit anti-c-FOS monoclonal antibody (Abcam, #ab214672), and 10 mg of mouse anti-NeuN monoclonal antibody (Encor Biotechnology, #MCA-1B7) followed by fluorescently conjugated secondaries in 1:2 primary : secondary molar ratios (Jackson ImmunoResearch). After active labeling, samples were incubated in EasyIndex (LifeCanvas Technologies) for refractive index matching (RI=1.52) and imaged at 3.6X with a SmartSPIM light sheet microscope (LifeCanvas Technologies).

Sample images were tile corrected, destriped and registered to the Allen Brain Atlas (Allen Institute: https://portal.brain-map.org/) using an automated process. A NeuN channel for each brain was registered to 8-20 atlas-aligned reference samples, using successive rigid, affine, and b-spline warping algorithms (SimpleElastix: https://simpleelastix.github.io/). An average alignment to the atlas was generated across all intermediate reference sample alignments to serve as the final atlas alignment value for the sample. Automated cell detection was performed using a custom convolutional neural network through the Tensorflow python package. The cell detection was performed by two networks in sequence. First, a fully-convolutional detection network (https://arxiv.org/abs/1605.06211v1) based on a U-Net architecture (https://arxiv.org/abs/1505.04597v1) was used to find possible positive locations. Second, a convolutional network using a ResNet architecture (https://arxiv.org/abs/1512.03385v1) was used to classify each location as positive or negative. Using the atlas registration, each cell location was projected onto the Allen Brain Atlas to count the number of cells for each atlas-defined region.

### RNA Scope in situ hybridization and quantification

Mice were euthanized using CO_2_ and DRGs were immediately dissected out and post-fixed in 4% PFA at 4°C for 24h, and then 30% sucrose for another 24h. Tissue was flash frozen on dry ice in OCT, sectioned at 12um onto Superfrost Plus Slides, and stored at -80 °C until processing using the Fresh Frozen RNA Scope protocol (ACD Bio). ACD Bio RNAscope Probes: Mm-Mrgprb4-C2 (435781-C2), Mm-Mrgprd (417921), and Mm-Mrgpra3-C3 (548161-C3). The fixation step in this protocol was skipped. Confocal images were obtained and quantified in Fiji.

### Stress paradigm

The protocol was based on the sub-threshold variable stress paradigm described by Hodes et al^27^. For the 3-day variable stress paradigm, stressors were administered in the following order: 100 random 0.45mA foot shocks for 1 hour, tail suspension for 1 hour, and restraint stress inside a 50ml falcon tube (decapicone for implanted animals) for one hour inside the home cage.

### Behavioral Assays Developmental milestones

Mouse pups underwent developmental milestone testing as described by Hill et al. ^68^, for PND 3-14. Assayed behaviors included surface righting (ability to flip over from back to abdomen <1 second), forelimb grasp (ability to grip a small wooden dowel >1 second), cliff aversion (reflexive movement away from a ledge), rooting (reflex to move head toward tip of cotton applicator stroked on cheek), and auditory startle (jumping reflex in response to handclap).

### Y Maze

Mice were habituated to the testing room for 1h before being placed in a y-shaped maze (15” x 5” x 3.5”/arm) with arms labeled A, B, and C. For training, one of the arms was closed off with a divider and the mouse was able to explore the 2 open arms for 15 minutes. The blocked arm and starting arm placement were recorded and counterbalanced. Mice were returned to their home cage for 1h, after which mice were placed back into the maze with all 3 arms open, and behavior was recorded for 5 minutes.

### Tape test

3cm of lab tape was placed on the backs of mice in 1000ml glass beakers and behavior was recorded for 5 minutes. Bouts and duration of licking, biting, sniffing, and shaking were scored.

### Sucrose Splash Test

This assay was performed under red light. Mice were habituated to the room for 1 hour before testing. Mice were sprayed on the back with 10% sucrose dissolved in water 3 times before being placed into an empty cage where behavior was recorded for 5 minutes. The amount of time spent grooming during the 5-minute period was hand scored by an observer blind to genotype. MrgprB4^Cre^; Rosa^DTA^ mice were single housed after this assay and Mrgprb4^Cre^; Rosa^hM3dq^ mice remained group housed.

### Novelty Suppressed Feeding

The assay was performed under red light. Mice were food restricted overnight ∼20h before testing. On the day of testing, mice were habituated to the room for 1 hour before being placed into a clean open field box (41 × 36 × 23cm). A single pellet of standard chow was placed in the center of the box. Latency to eat was scored up to 10 minutes.

### Sucrose Consumption

Mice were given 2 50ml bottles in their home cage: one containing plain water and other containing 1% sucrose. After 24 hours, the 2 bottles were weighed to calculate consumption, and the position of the bottles was switched for an additional 24 hours. The amount consumed over the 2 days was averaged.

### Forced Swim Test

Animals were habituated to the testing room for 1 hour before testing. Mice were placed in 4L plastic beakers containing water at ∼25°C for 6 minutes. Behavior was recorded for 6 minutes and behavior for last 4 minutes of the trial were scored by an observer blind to genotype.

### Open Field Test

Mice were acclimated to the testing room for 1 hour before being placed into open field arena (42 × 42 × 40cm) for 30 minutes. Behavior was recorded and distance traveled was measured using Noldus Ethovision software.

### Electrode Implantation

Mice were bred from Rosa-DTA cross with B4-Cre mice, genotyped and weaned at 21 days of age. At 6-7 weeks old, male and female, wild-type and Cre positive mice were anesthetized with isoflurane, placed in a stereotaxic device, and underwent craniotomy to implant electrodes. Metal ground screws were secured to the cranium above the olfactory bulb and cerebellum. Tungsten microwires (California Fine Wires) were implanted in the prelimbic and infralimbic cortices (PrL, IL), basolateral and central amygdala (BLA, CeA), Nucleus Accumbens (NAc) and ventral tegmental area (VTA) as previously described^41^. At the conclusion of behavioral and neurophysiological experiments, animals were euthanized via cardiac perfusion with 4% paraformaldehyde and histological analysis and CT scanning of electrode placement was performed to confirm recording sites.

### Neurophysiological Recordings

Prior to recordings, animals underwent the subchronic variable stress paradigm for one week. Mice were connected to Mu headstages (Blackrock Microsystems Inc., UT) under isoflurane anesthesia 60 minutes prior to recording sessions. During recordings, animals were placed in a homecage setting for 5 minutes. Animals were then sprayed on their backs with a 10% sucrose solution (as described in Sucrose Splash Test) three times then recorded for 5 minutes. Post-sucrose splash test, animals were recorded again for 5 minutes. Neural activity was recorded at 30kHz using the Cereplex Direct acquisition system (Blackrock Microsystems Inc., UT). Local field potentials (LFPs) were bandpass filtered at 0.5–250Hz and stored at 1000Hz. Recordings were referenced to a ground wire connected to ground screws in the skull.

### Directionality Analysis, Frequency Band Selection, and Power

Signal directionality between brain regions was estimated using a temporal lag approach as previously described^45^. Briefly, LFP data recorded from B4^Cre^; Rosa^DTA^ and +/+;Rosa^DTA^ during homecage recordings was filtered using butterworth bandpass filters to isolate LFP oscillations within a 1Hz window using a 1Hz step in the 1-50Hz range. The Hilbert transform was used to determine the instantaneous phase of the filtered LFPs. The instantaneous phase offset time series was calculated for each LFP pair (ϕArea2 -ϕArea1)t. The mean resultant length (MRL) for the phase offset time series was calculated, representing the phase coherence between pairs of LFPs. Temporal offsets were introduced between each pair of LFPs ranging from -250ms to 250ms in 2 ms steps and then the phase coherence was recalculated between each LFP pair. The offset at which the two LFPs optimally phase synchronized is plotted for each frequency +/-sem.

In order to hone in on the exact frequency bands at which each brain region was leading or lagging, we used a fuse lasso approach. Lasso (<u>L</u>east <u>a</u>bsolute <u>s</u>hrinkage and <u>s</u>election <u>o</u>perator), is a method of model parameter selection and regularization. The lasso method minimizes the least squares loss over a set of parameters in a linear regression problem by imposing the condition that the sum of the absolute values of the parameter estimates (that is, the 𝓁_1_ norm of the parameters) is less than some value^69^. This method usually results in some parameter estimates being zeroed out, in essence selecting the important covariates that are more predictive of the response than those with zeroed out regression coefficients. The fused lasso generalizes the lasso in the sense that it accounts for temporal or spatial dependence amongst covariates by further imposing a condition that the sum of the absolute values of the lag one differences between subsequent parameter estimates be less than some value^70^. The problem of modeling the response under fused lasso then amounts to finding a set of parameter estimates 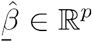 as a solution of

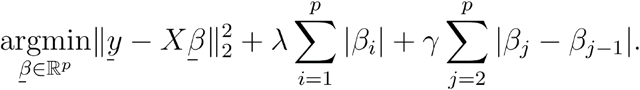

The parameters λ and are *γ* positive tuning parameters. The solution of the optimization problem in (1) depends on the chosen value of λ and *γ*, that is 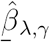, which are chosen via cross validation in such a way that the test mean squared error is minimized.

We used fused lasso for approximating the significant frequency bands within the 1*Hz* to 50*Hz* range obtained from the the phase offsets data for eight different mice and for each pair of brain regions from the set: BLA, CeA, IL, NAc, PrL, VHipp and VTA. In our case, our hypothesis is that a small number of frequencies are active during the measurements and an active frequency implies that neighboring frequencies are also active. Our setup follows from the application of fused lasso for signal approximation^71^. In order to use fused lasso to select significant frequency bands, we first computed the mean over all observations for each pair of brain regions and for each mouse. This resulted in 168 observations (8 mice times 21 brain region pairs) each with 50 elements corresponding to the mean phase offset at each frequency from 1*Hz* to 50*Hz*. This 168 × 50 matrix was then reshaped into a 400 × 21 response matrix **Y** in which each column contained the means across frequencies for each mouse for a particular pair of brain regions. The design matrix **X** was comprised of eight 50 × 50 identity matrices stacked on top of each other (one for each mouse) resulting in a 400 × 50 matrix. The 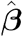 matrix of parameter estimates was a 50 × 21 matrix for which each column contained the model estimates for the phase offsets over the 1*Hz* to 50*Hz* range for a given pair of brain regions. We then used the genlasso library^72^ in R to obtain estimates 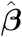 under the fused lasso model with the tuning parameters λ and *γ* set to 3 and 0.5 respectively. This resulted in 21 different vectors (the columns of 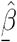) with bands of non-zero values corresponding to significant phase offset frequencies and zeros indicating non-significant frequencies for each pair of brain regions.

### Electrode Placement Identification

Two primary methods were used for checking electrode placements: computerized tomography (CT) verification of electrode placement using a 3D configuration and histology. For CT scans, implanted animals were perfused with 4% PFA and submerged in 4% PFA overnight with electrodes still intact. The samples were then placed in 7.5% Lugol’s solution for 24 hours before microCT imaging. Samples were imaged by the Small Animal Imaging Core at the Iowa Institute of Biomedical Imaging. All animals utilized in the neurophysiological recording experiments were perfused with 4% PFA following recordings. Brains were stored in 30% sucrose overnight and frozen in OCT compound on dry ice and stored at -80°. Brains were sectioned into 35μm slices and washed in PBS following standard methods and stained with red Nissl (ThermoFisher, Waltham, MA). Slices were imaged using a Leica fluorescence microscope at 10x magnification. Following imaging, brains were aligned with the Paxinos & Franklin brain atlas for verification of electrode placements. Electrode wires are outlined in blue for each target brain region.

### Corticosterone ELISA

The mild acute stress used for corticosterone measurements prior to the variable stress paradigm involved removal from housing room, brief restraint, and heating of the tail using a heating pad. Comparatively, baseline measurements were obtained after collecting blood in the housing room without restraint. Tail nick bleeds were taken at the described timepoints, and blood was placed into EDTA-treated tubes. Tubes were spun in a centrifuge at 5000 rpm for 10 minutes and plasma was collected. Plasma aliquots were stored at -20°C until analysis. CORT levels were measured using a commercially available ELISA kit (Abcam) according to the manufacturers’ instructions.

### Experimental Design and Statistical Analysis

All statistical analyses were performed using Graph Pad Prism 9.0 software. Statistical significance was set at p<0.05. Samples that varied >2SDs from the mean were excluded.

## Acknowledgements

We thank members of the Abdus-Saboor, Hultman, and Blendy labs for thoughtful comments and feedback on this work and manuscript. We thank Alli Jimenez and Radha Velamuri for technical support. MDS thanks members of her thesis committee for helpful feedback on this work. MDS is supported by NIH NRSA grants from NCCIH. IAS and lab members acknowledge support from startup funds provided by the University of Pennsylvania and Columbia University, National Institute of Health grant NIH/NIDCR R00-DE026807, and fellowships from the Rita Allen Foundation and Alfred P. Sloan Foundation. RH thanks Iowa Neuroscience Institute and Carver Trust.

---

**Supplemental Figure 1.**
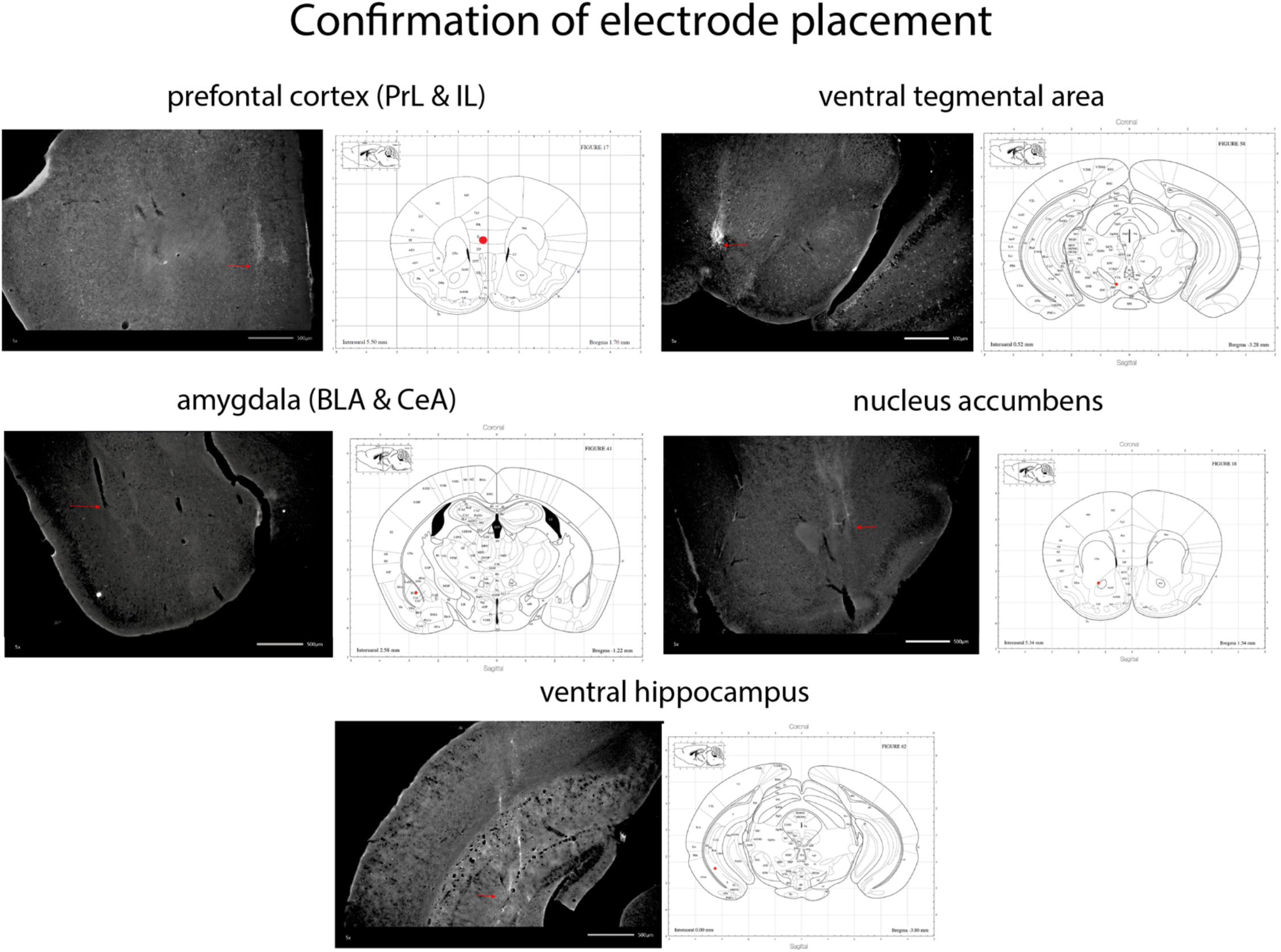
Histological confirmation of electrode placement. Red arrow indicates site of electrode tip. Red dot indicates target brain region in mouse brain atlas.

**Supplemental Figure 2.**
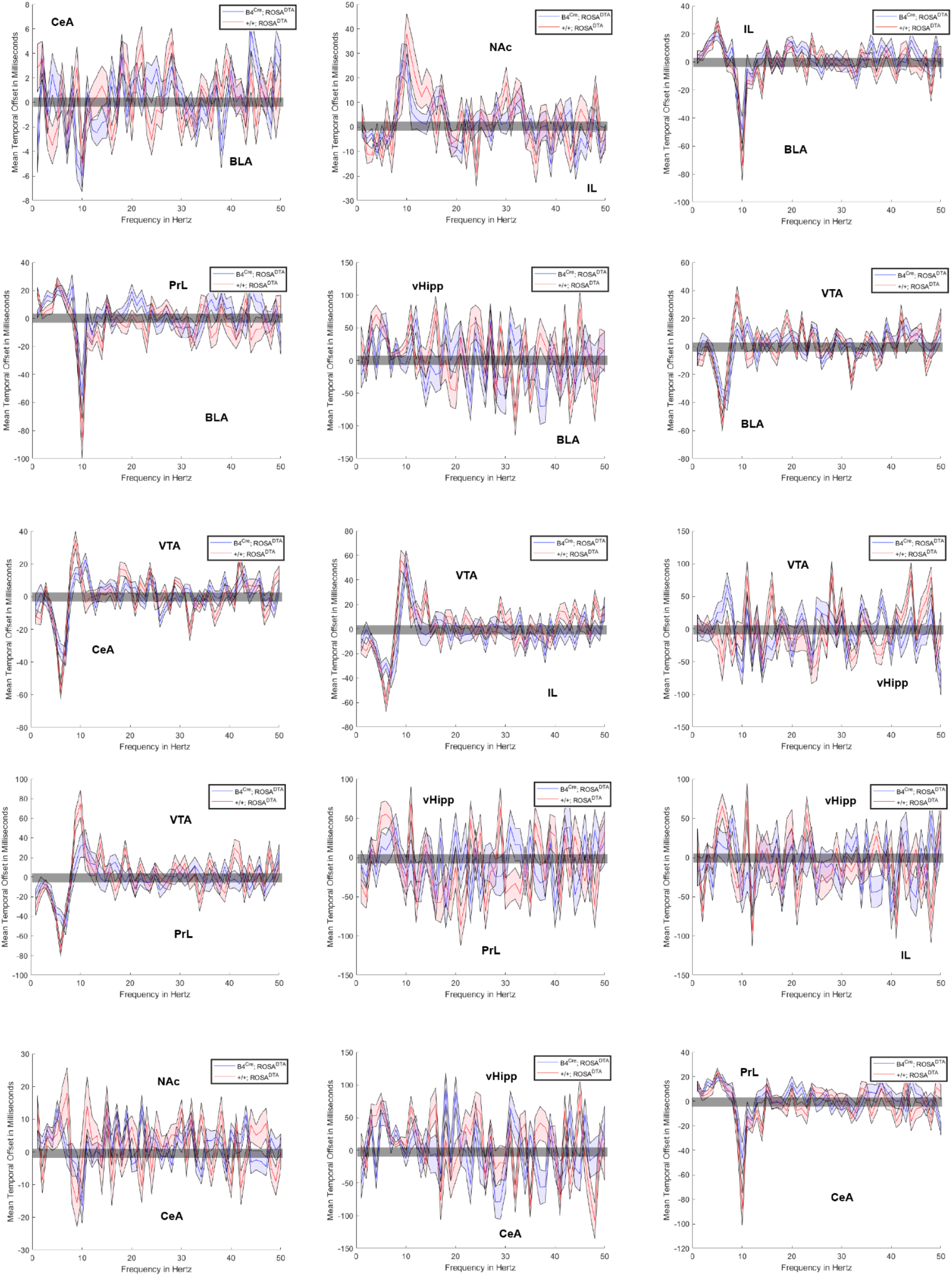
Directionality plots for brain region pairs. Temporal offset above zero indicates that the brain region listed above zero is leading. Brain region listed below the 0 line is leading when temporal offset is below zero.

**Supplemental Figure 3.**
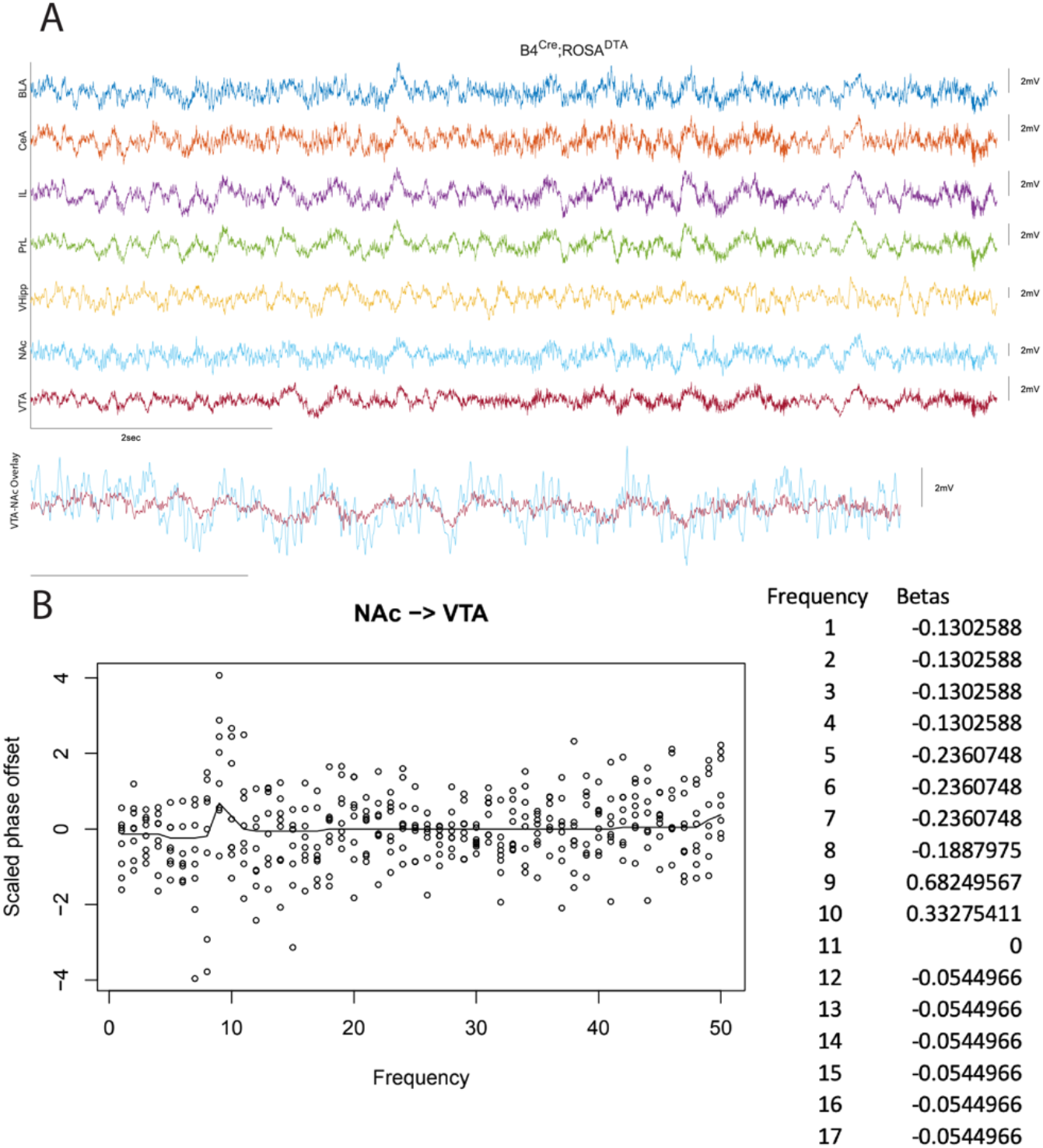
Lasso-based methods were used to identify frequency bands of interest. **A**, Representative LFP traces from a Mrgprb4-albated animal. **B**, Frequencies in the 1-10Hz range had a scaled phase offset furthest from zero.

